# Eyes-closed resting state EEG reveals clearer and more stable group differences between autistic and neurotypical individuals than eyes-open resting state EEG

**DOI:** 10.1101/2025.01.13.632659

**Authors:** Wenyi Xiao, Myles Jones, Elizabeth Milne, Adam J.O Dede

## Abstract

**Background:** Resting-state electroencephalography (rs-EEG) has been widely used to explore neural dynamics in Autism Spectrum Condition (ASC). However, inconsistencies in findings across studies remain a challenge, partly due to variations in brain state, such as eye conditions (eyes-open vs. eyes-closed). This study aims to examine rs-EEG differences between ASC and neurotypical (NT) participants, focusing on the influence of eye condition.

**Methods:** A total of 300 participants (126 ASC) were included. Rs-EEG data were analysed across eyes-open, eyes-closed, and difference between eye conditions, with 726 variables assessed per participant. Linear regression and effect size (η^2^*_partial_*) were used to identify group differences, complemented by cluster-based permutation testing and bootstrapped split-half validation for reliability.

**Results:** Group differences were most pronounced in the eyes-closed condition, particularly for relative power and multiscale entropy (MSE). Compared to neurotypical participants, ASC participants exhibited reduced frontal coarse-scale MSE, increased delta power, and decreased alpha power, suggesting altered local-global neural dynamics. Cross-validation revealed greater reliability of effects in the eyes-closed condition compared to eyes-open or difference between eye conditions.

**Conclusions:** Eye condition plays a critical role in detecting rs-EEG differences between ASC and NT groups, with the eyes-closed condition yielding more consistent and pronounced effects. These findings highlight the importance of controlling brain state in rs-EEG studies and suggest that integrating eye condition effects with other biomarkers may improve identification of neural differences associated with ASC.

## 1. INTRODUCTION

Autism Spectrum Condition (ASC) is a neurodevelopmental and heterogeneous condition. Resting-state electroencephalography (rs-EEG) has emerged as a valuable tool for investigating the neural underpinnings of ASC, offering a non-invasive and easily-collected method to measure brain activity (Jeste, Frohlich, & Loo, 2015). A common research strategy involves comparing rs-EEG data from individuals with ASC to that of neurotypical (NT) individuals to identify neurobiological differences that could contribute to ASC diagnosis.

Despite extensive research, identifying consistent neurobiological differences between ASC and NT individuals remains challenging. Conflicting findings have been reported, including increases in absolute alpha power (Cornew et al., 2012) and decreases in absolute alpha power (Wang et al., 2013). Similarly, studies on multiscale entropy (MSE), a measure of neural signal complexity, have reported reductions in ASC (Catarino et al., 2011; Milne et al., 2019), but it remains underexplored and lacks sufficient replication studies (Ribeiro & da Silva Filho, 2023). Conflicting findings have been reported for other variables as well (Dickinson et al., (2018) O’Reilly et al. (2017) Garcés et al. (2022)). These inconsistencies may result from overlapping patterns of brain activity between the groups, with only small or inconsistent effects observed across metrics (Garcés et al., 2022), emphasising the need for larger cohorts and validation.

One reason for conflicting results may be the heterogeneity of ASC itself combined with small sample sizes. In our previous work which compared resting-state EEG data obtained during eyes-open rest, utilising a sample of 776 participants, we found that many of the differences previously reported between ASC and NT individuals generate small effect sizes and fail to replicate using split-test validation methods (Dede et al., 2023). Notably, our analysis also revealed that larger effect sizes tend to emerge in smaller sample sizes, but these effects are less likely to replicate. This issue is acute given that 85% of studies in a recent meta-analysis investigating differences in spectral power and connectivity between neurotypical and autistic people included fewer than 50 ASC participants, highlighting a widespread limitation in sample size (Neo et al. 2023).

Inconsistencies in rs-EEG findings may also stem from variations in the resting-state paradigm. A particularly important factor is eye condition, as brain activity differs significantly depending on whether participants’ eyes are open or closed (Berger 1929). Neo et al. (2023) highlighted the importance of this variable, showing that effect sizes varied between studies utilizing eyes open versus eyes closed rs-EEG. Additionally, the mental state of participants in ASC studies is often under-controlled, which may further contribute to variability (DiStefano et al., 2019). Despite these observations, the role of eye condition in modulating rs-EEG group differences between ASC and NT individuals remains underexplored. To date, only one study has examined this contrast within subjects, focusing solely on alpha power (Bellato et al., 2020).

In the present study, we computed the same variables as in our previous report and drew from the same database of participants (Dede et al., 2023). However, we constrained our analysis to a subset of 300 participants for whom both eyes-open and eyes-closed rs-EEG data were available. In this way, it was possible to evaluate whether changes in brain state interacted with autism diagnosis to reveal group differences that had not been apparent when examining eyes open data alone. Our investigation centres on three key questions: (1) how eye conditions impact the detection of ASC group differences in rs-EEG metrics, (2) which rs-EEG measures most effectively predict ASC diagnosis under different conditions, and (3) the reliability of these group differences through cross-validation.

## 2. METHODS

This project was pre-registered (Xiao & Dede, 2023), however alterations were made to the analysis plan in order to better summarise the data and increase statistical rigour. For clarity of exposition, we do not note the differences here, but the interested reader is invited to compare the present methods to our pre-registration.

Due to the large dataset samples, analysis depended on high power computing (HPC) resources provided by the University of Sheffield. For details on code optimization, see our earlier report (Dede et al., 2023). Code and summary EEG variables are available in the GitHub repository: https://github.com/adede1988/SheffieldAutismBiomarkers.

### 2.1 Data

No new data were collected for this project. We selected the subset of participants from our previous study (Dede et al., 2023) for whom both eyes open and eyes closed data were available. This resulted in only data from the Autism Center for Excellence (ACE) project Multimodal Developmental Neurogenetics of Females with ASD being used (Neuhaus et al. 2021). De-identified data were obtained from the National Database of Autism Research (NDAR study #2021).

Initially, the dataset consisted of 336 participants. After excluding participants for whom more than 50% of channels were removed due to noise, or who were missing age information, the dataset was reduced to 333 participants with eyes-open data and 305 with eyes-closed data. Only participants with data available for both eyes-open and eyes-closed conditions were retained, resulting in 300 participants in the final dataset, including 126 ASC participants (Table 1).

**Table 1.** Demographic characteristics of participants and relevant EEG parameters.

| Characteristic | ASC | NT |
| --- | --- | --- |
| Total number (women) | 126 (53) | 174 (89) |
| Age in months | 153 ± 33 | 156 ± 35 |
| ADOS | 11.64 ± 2.62 | / |
| IQ (DAS GCA) | 107 ± 21 | 113 ± 15 |
| Original number of EEG channels | 125 ± 0 | 125 ± 0 |
| Final number of EEG channels | 125 ± 1 | 125 ± 1 |
| Original number of EEG epochs | 69 ± 26 | 69 ± 26 |
| Final number of EEG epochs | 65 ± 28 | 59 ± 28 |

*DAS GCA = Differential Ability Scales General Conceptual Ability.**Mean values are presented with ' ± ' standard deviation values.***Note:** No ADOS data was collected for NT participants.

Autistic participants were assessed using the Autism Diagnostic Observation Schedule (ADOS), with the module tailored to their age and language ability. Neurotypical participants were screened using either the ADOS or the professional judgement of qualified clinicians.

Participants were classified as ASC (ADOS > 7) or neurotypical (NT) based on standardised ADOS scores or clinical evaluation (Lord et al., 2012).

### 2.2 Preprocessing

EEG data were preprocessed by the original data collection team. Data were high pass filtered at.1 Hz, low pass filtered at 100Hz, and notch filtered at 60 Hz. Channels with high impedances were removed, and the data were segmented into 2.048-second epochs. The difference between the maximum and minimum voltage in a moving 80 ms window was calculated for each channel and epoch. Epochs with at least one 100 LV deflection were flagged as potentially noisy. A channel was deemed ‘bad’ if it crossed the 100 LV threshold in 50% of trials. ‘Bad’ channels were removed. A trial was deemed ‘bad’ if 25% of the remaining channels exhibited threshold crossings. ‘Bad trials’ were removed. As preprocessing was conducted externally, we could not assess noise differences between groups. However, a previous report using these data found that ASC participants exhibited more noisy data (Neuhaus et al., 2021).

To ensure a standard electrode montage, simplify analysis, and increase signal to noise ratio, all data were referenced to an average reference and interpolated to a standard 32-channel montage. Finally, a Laplacian transform was applied to estimate the current source density of the data. The Laplacian transformed data were used for the calculation of inter-site phase clustering (ISPC).

### 2.3 rs-EEG metrics calculation

We calculated power spectra, 1/f slope (the slope of the power spectral density), peak alpha frequency (PAF), phase-amplitude coupling (PAC), multiscale sample entropy (MSE), and inter-site phase clustering (ISPC) using established methods. For details of the calculations, refer to our previous study (Dede et al., 2023). Power spectra were computed by averaging the narrowband-filtered power time series across epochs, using Gaussian convolution for filtering in the frequency domain. The 1/f slope was estimated via linear fitting, following Voytek et al. (2015). PAF was calculated as the mean of the best-fitting Gaussian within a 6 to 14 Hz range (Dickinson et al., 2018). PAC was quantified using methods described by Tort et al. (2010), with frequency bands defined as per Peck et al. (2022). MSE was derived by adjusting data granularity and calculating entropy across scales, based on methods from Costa et al. (2005) and Busa & van Emmerik (2016). ISPC was computed on Laplacian-transformed data using standard protocols (Cohen, 2014).

These metrics were previously calculated and archived in the Sheffield Autism Biomarkers repository, with extracted component values available for reference (Dede et al., 2023).

### 2.4 Statistical evaluation of group differences across all variables

All measures were regressed onto the age, sex and autism diagnosis variables.

Dependent measure ∼ age + sex + autismDiagnosis

We conducted a total of 2,178 statistical tests by performing 726 tests for each of three conditions (eye open, eye closed, and difference between eye conditions).

The coefficients were estimated in the R statistical environment (R core team, 2021) using linear regression (*lm* function). For each test, η^2^*_partial_* associated with each independent variable (age, sex and diagnosis) was calculated to measure effect size. In all cases, type III sums of squares were used to calculate η^2^*_partial_*, which means that all effect size values reflect the effect of a predictor assuming that all other predictors were added into the model before it. Due to the large number of tests performed and the large sample size, it was expected that many would yield p-values below.05. Thus, the present analysis focused on η^2^*_partial_* instead of p-value because it is not subject to the same multiple comparison issue, and where a significant p-value observed with a large sample size is sometimes associated with a negligible effect, assessing effect size directly circumvents this issue. In addition, this analysis served as a data-driven way to select a subset of variables for further exploration in our fine-grained analysis (Section 2.6).

### 2.5 Aggregation of rs-EEG measures

All rs-EEG measures were calculated per channel, except for ISPC, which was calculated for channel pairs. However, due to correlated signals at adjacent electrodes and the impracticality of analysing all values, measures were aggregated into predefined electrode groupings and frequency bands, as used in prior work (Dede et al., 2023). Electrode groupings included 13 scalp regions and 5 asymmetry-sensitive comparisons contrasting signal in one scalp region compared to another (e.g. medial minus lateral electrodes).

For power spectrum comparisons, spectra from each channel were averaged in 6 frequency bands: δ (2-4 Hz), θ (4-8 Hz), α (8-14 Hz), L (14-30 Hz), Llow (30-50 Hz), and Lhigh (50-80 Hz). These band-averaged power values were used for all regional and asymmetry comparisons and repeated for raw, log-transformed, and relative power measures, resulting in 324 dependent measures (18 comparisons X 6 frequencies X 3 measures).

The 1/f trend slope was calculated using both log-transformed and relative power values, with regional and asymmetry comparisons yielding 36 dependent measures (18 comparisons × 2 measures). Similarly, for peak alpha frequency (PAF), regional and asymmetry comparisons were computed for values derived from both log-transformed and relative power data, resulting in another 36 dependent measures (18 comparisons × 2 measures).

Phase-amplitude coupling (PAC) involved dividing low frequencies into δ (2-4 Hz), θ (4-8 Hz), α (8-14 Hz), β (14-20 Hz), and high frequencies were divided into β (20-32 Hz; note when β was compared to β, 24 Hz was used as the low cut off for the higher frequency), γlow (32-52 Hz), and γhigh (52-100 Hz). This created 12 frequency band pairs for which PAC was averaged across channels. Regional and asymmetry comparisons were calculated z-scored PAC values relative to shuffle-generated null distributions, resulting in 216 dependent measures (18 comparisons × 12 frequency pairs).

For multiscale entropy (MSE), scale ranges were categorised into all scales (1–20), fine scales (1–7), medium scales (8–13), and coarse scales (14–20). MSE values were averaged within these ranges and used for all regional and asymmetry comparisons, producing 72 dependent measures (18 comparisons × 4 scale ranges).

Inter-site phase clustering (ISPC) was computed within the same six frequency bands used for power spectra. The average ISPC was calculated for electrode pairs specified in the five asymmetry comparisons, as regional comparisons were not applicable to ISPC. Long-and short-distance connectivity were also calculated by averaging ISPC across channel pairs with inter-electrode Euclidean distances greater or less than the median, respectively. This yielded 42 dependent measures (7 comparisons × 6 frequency bands).

In total, 726 dependent measures (324 power spectra + 36 1/f slope + 36 PAF + 216 PAC + 72 MSE + 42 ISPC) were analysed across three conditions: eyes open, eyes closed, and the difference between the two. This resulted in 2,178 dependent measures.

### 2.6 Assessing effect size Stability and Predictive Power of Independent Variables

To assess the stability of the identified coefficients, we employed bootstrapped split-half cross-validation. This process involved several steps: 1) randomly dividing the data into two halves, ensuring that both halves maintained equal proportions of participants in each diagnostic group, as well as balanced distributions of age and sex; 2) refitting the models to each data split and recalculating the effect sizes; 3) repeating this process across 150 random data splits and determining the proportion of splits where both halves yielded η^2^*_partial_* values above.035 for each variable. This proportion was termed the replication rate. We focused on variables with replication rates of at least.64, corresponding to the convention of.80 statistical power. In essence, if tests are considered reliable when they detect an effect with 80% probability, a replication rate of.64 is acceptable when repeating the test. The.035 threshold for η^2^*_partial_* was selected because it falls between the standard definitions of small and moderate effect sizes (Cohen, 1988), and effects below this threshold are unlikely to be of practical significance.

### 2.7 Fine-grained analysis

Of 105 EEG measures that were predicted for diagnosis with η^2^*_partial_* >.035, 67 of them were either relative power or entropy measures (Figure 1, Supplementary Table 1). Chi-squared tests confirmed that these two types—relative power and entropy—were significantly overrepresented compared to other EEG measure types (Section 3.2). This finding motivated a more detailed exploratory analysis of these variables across the entire frequency and scale spectra at the single-channel level. A series of two-tailed cluster permutation tests were performed using threshold-free cluster enhancement (TFCE) (Friston et al., 1996;Smith & Nichols, 2009; Maris & Oostenveld, 2007;Mensen & Khatami, 2013). TFCE was applied across frequencies and scales to identify significant clusters, rather than running separate analyses at each point. For the TFCE analysis, signal intensity (H) was set to 2, and spatial distribution (E) was set to 0.66, following the guidelines from Mensen and Khatami (2013). Hypothesis testing utilised a Monte Carlo approach with 5,000 random permutations. These analyses were conducted using the ept_TFCE function from the TFCE toolbox (https://github.com/Mensen/ept_TFCE-matlab).

**Figure 1.**
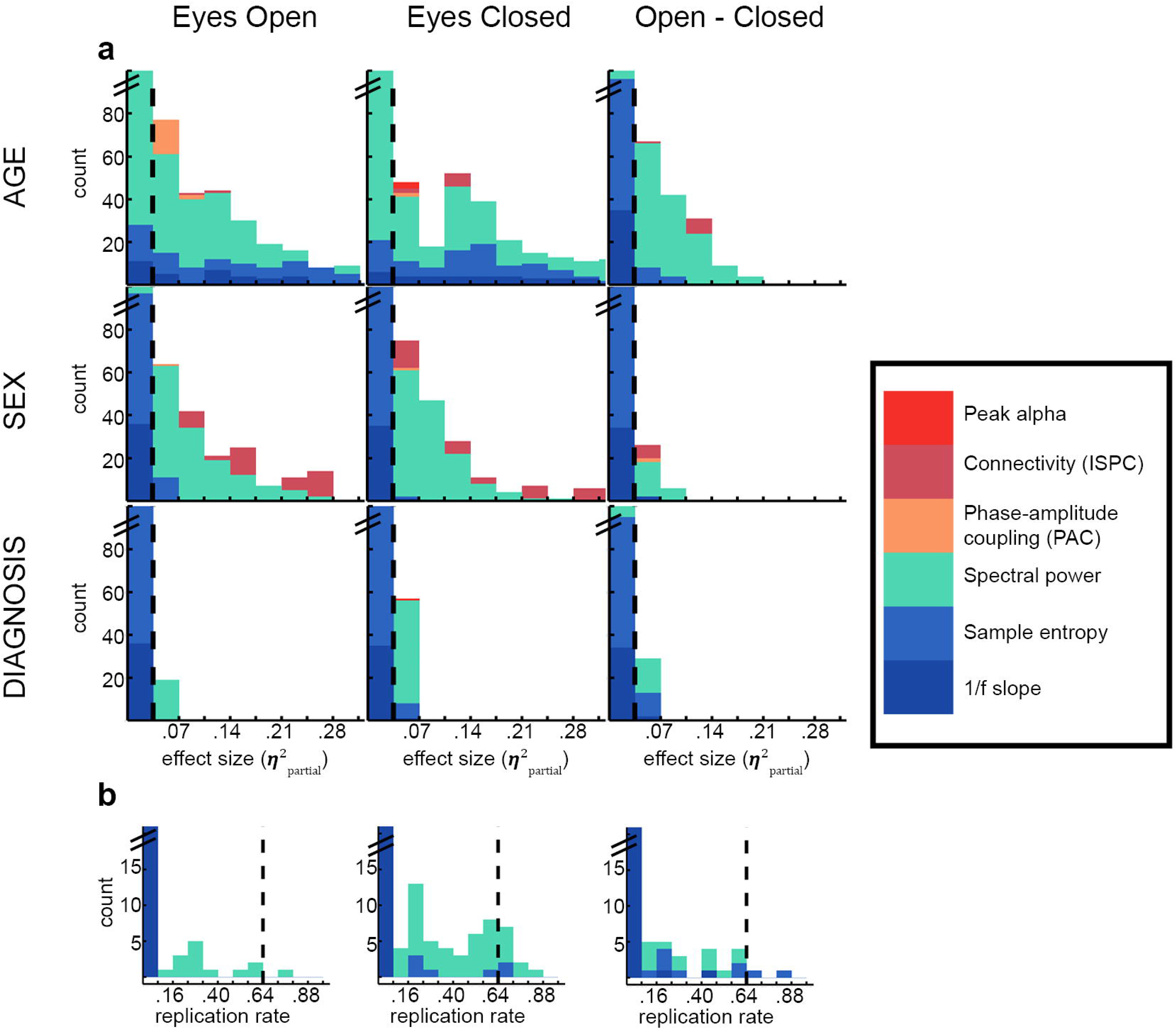
**a**.Effect sizes of age, sex, and diagnosis as predictors of resting-state EEG features across different eye conditions (eyes open, eyes closed, and open-closed difference). Age and sex were strong predictors across all conditions, while diagnosis showed limited predictive power, with higher effect sizes in the eyes-closed condition. Among the EEG metrics, power and sample entropy demonstrated greater effect sizes for diagnosis, particularly in the eyes-closed condition.The histograms display the distribution of effect sizes for the independent variables (rows) across the specified eye conditions (columns). The colour coding represents the different EEG-dependent variables (see legend). Y axes are truncated to emphasise larger effect sizes. Dashed vertical lines at.035 indicate the threshold used to separate meaningful effect sizes from noise. Supplementary Table 1 details the counts of significant dependent variables predicted by the independent variable ‘diagnosis’ in each eye condition (η^2^*_partial_* >0.035). **b.**The eyes-closed condition demonstrates more stable variables compared to the eyes-open condition, as well as the differences between the two conditions. Histograms illustrate the reliability rates for effect sizes greater than 0.035 (η^2^*_partial_* >0.035). These reliability rates, ranging from 0 to 1, were computed by comparing corresponding effect sizes in independently analysed training and test datasets, with higher values indicating greater stability. Colours denote the type of dependent variable (refer to the legend). The vertical dashed lines at 0.064 mark the threshold distinguishing stable effect sizes from unstable ones. Table 2 provides counts of dependent variables that were accurately predicted by the independent variable “diagnosis” in each eye condition. Well-predicted variables meet both the effect size and stability thresholds.

## 3. RESULTS

### 3.1 Quantitative search to identify diagnosis group difference in both eye conditions

There were examples of all EEG measure types that varied as a function of age, sex and/or diagnosis. Figure 1 displays histograms of the effect sizes associated with different predictors in all three conditions. Visual inspection of Figure 1 indicates that both age and sex exhibited strong prediction of EEG dynamics as reflected by a large number of η^2^*_partial_* values above.035.

By contrast, there was a paucity of variables that could be well-predicted by diagnosis, which extends our prior results to the eyes closed condition (Dede et al., 2023). Note that for the eyes open condition, this result is drawn from a subset of the same analysis presented in our previous report.

### 3.2 Distribution of well-predicted variables by type and condition

Among 2,178 model fits across three resting state conditions, only 105 dependent measures varied as a function of ASC diagnosis ( η^2^*_partial_* >.035; Figure 1 and Supplementary Table 1). In other words, only 5% of EEG metrics varied with ASC diagnosis. A chi-squared test revealed that more EEG variables were predicted by diagnosis in the eyes closed condition than in the other two conditions (x^2^(2) = 22.17, p <.001). In fact, 57/105 or 54.3% of meaningful effects were detected in the eyes closed condition.

Next, we explored which aspects of EEG dynamics were best predicted by ASC diagnosis. A chi-squared test examined whether the 105 meaningfully-predicted EEG variables were evenly distributed across 8 types of rs-EEG metrics: Power metrics (raw, relative, log), MSE, PAC, ISPC, PAF, or aperiodic slope. Results revealed that relative power variables yielded more meaningful effects than would have been expected by chance (49/105; 46.7%; x^2^(1) = 83.80, p <.001). Similarly, more MSE variables were predicted by diagnosis than would have been expected by chance (18/105; 17.1%; x^2^(1) = 6.47, p = 0.011). The remaining variables were measures of Log Power, raw power, aperiodic slope and PAF, but these variables were not represented with frequencies greater than would be expected by chance.

These findings highlight the importance of measure type in EEG associations with ASC diagnosis, with multiscale sample entropy and relative power showing the strongest variation. Relative power and multiscale sample entropy in the eyes-closed condition emerged as key indicators. However, these results were based on averages across electrodes and within frequency bands/scale factor ranges, and they were not cross-validated. Further cross validation and then fine-grained analysis at the single electrode level and without averaging across either scale factors or frequencies was also conducted (Section 2.6).

### 3.3 Cross Validation Results

Figure 1b illustrates the replication rate of diagnosis effects. The replication rate was the proportion of the time across repeated bootstrap samples that both random halves of the data exhibited a replication of the same effect. Replication rate was investigated separately within each eye condition. EEG measures with both a diagnosis group difference of η^2^*_partial_* >.035 and a replication rate greater than.64 were considered well-predicted. 13 variables were identified as being well-predicted (Table 2). Of variables that were identified as simply yielding η^2^*_partial_* >.035, 1 of 726 in the eyes-open condition had replication rates above.64. The corresponding values for the eyes-closed and difference between eye conditions were: 10 of 726 variables and 2 of 726 variables, respectively. The eyes-closed condition had more well-predicted variables than the other conditions (x^2^(2) = 11.23, p = 0.0036).

**Table 2.**
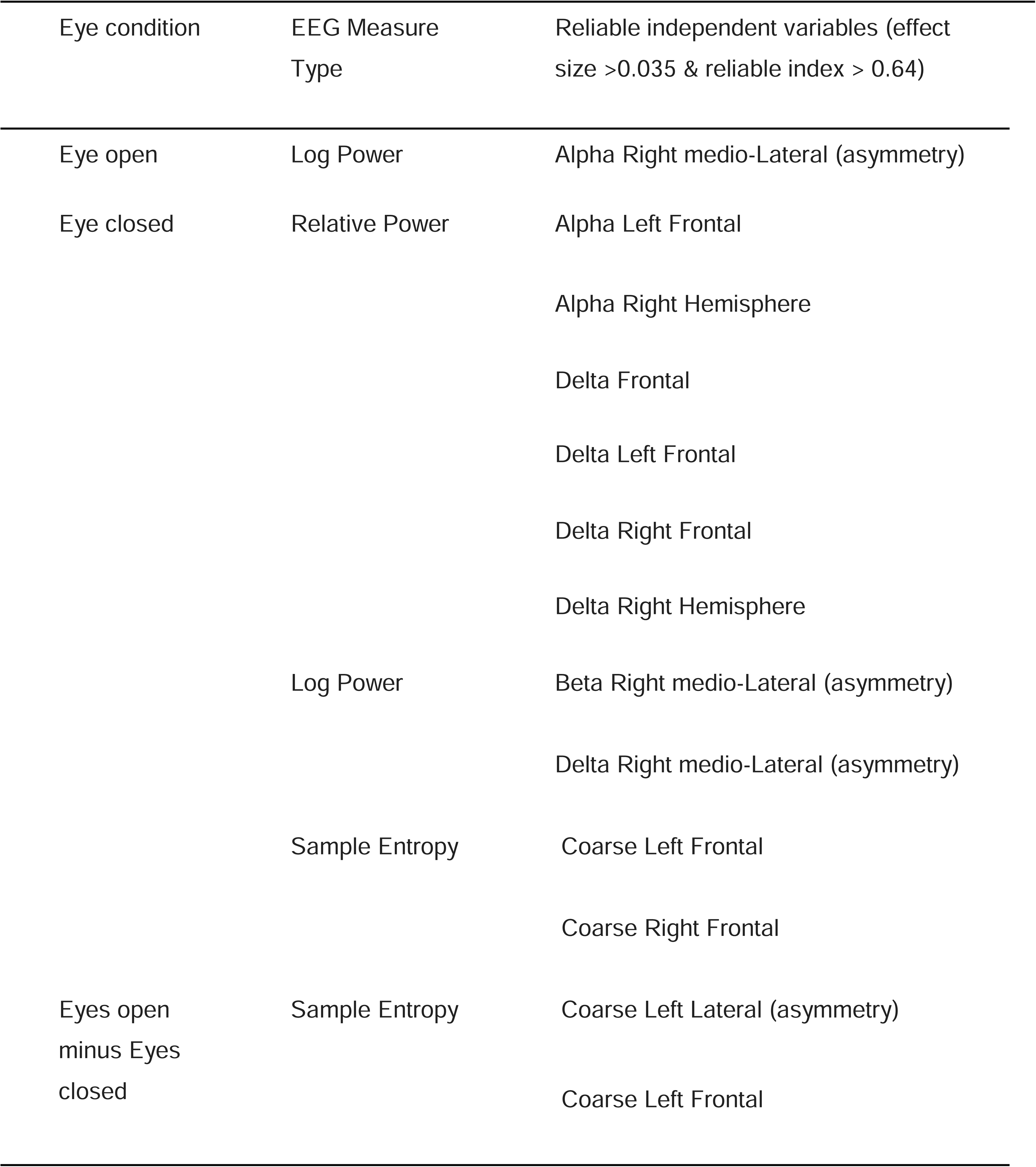
Reliable Univariate Independent Variables Predicting Diagnosis Across Eye Conditions (Eyes-Open, Eyes-Closed, and Eyes-Open Minus Eyes-Closed)

### 3.4 Fine-grained Analysis of Relative Power and Multiscale Entropy

Individual data points for relative power and MSE, averaged across brain regions, within each diagnosis group were visualised in Figure 2. Visual examination of these plots reveals that the indices align with expectations. All participants exhibited higher sample entropy at more coarse relative to fine scales. The alpha peak is more prominent when the eyes are closed, and the classic 1/f scaling of the power spectrum is evident.

**Figure 2.**
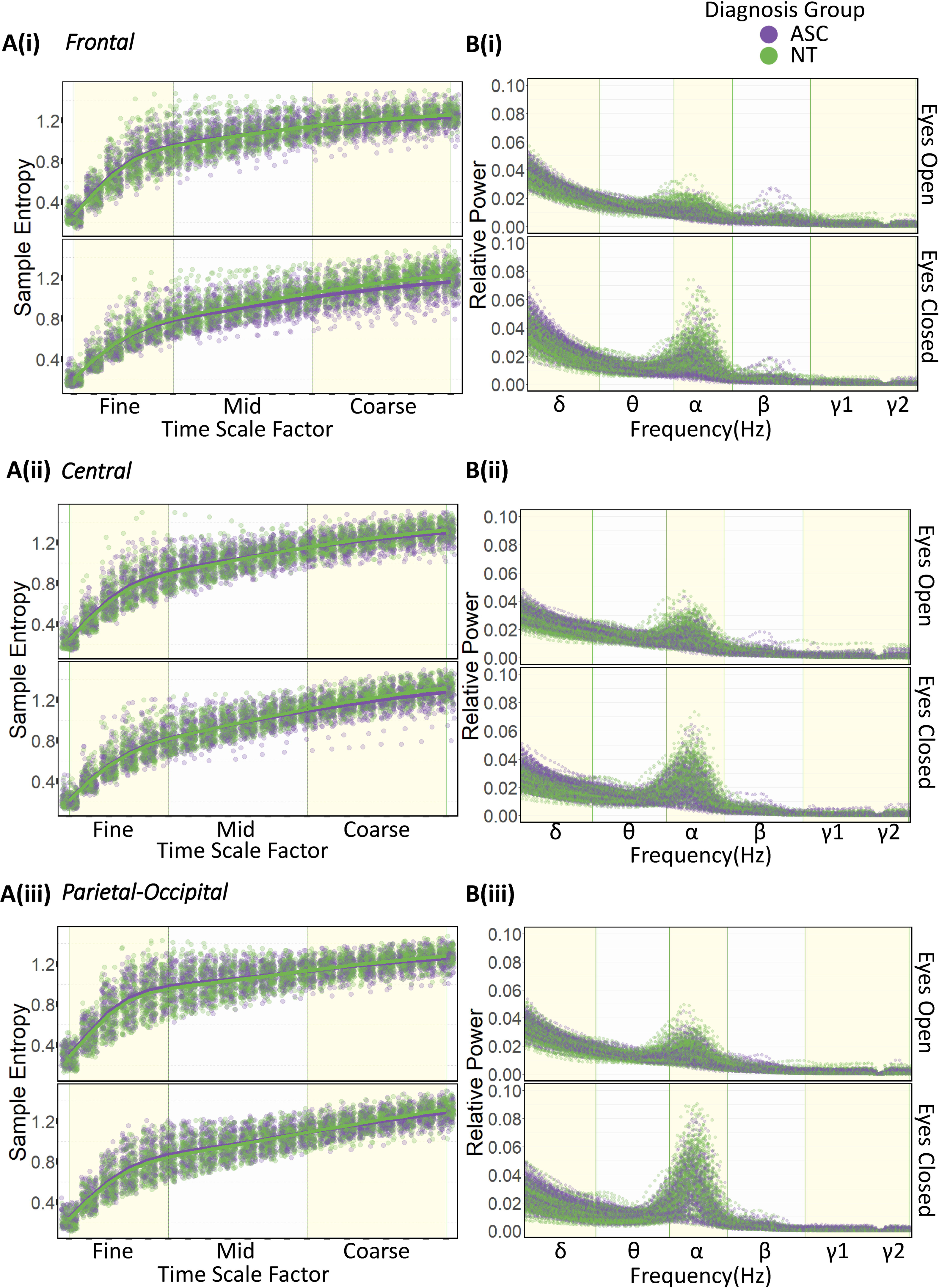
Panels A(i), A(ii), and A(iii) display Sample Entropy across time scale factors (fine, mid, coarse) for frontal, central, and parietal-occipital regions, respectively, in ASC and NTgroups. Panels B(i), B(ii), and B(iii) illustrate Relative Power across frequency bands (δ, θ, α, β, γ1, γ2) for the same brain regions under eyes-open (top rows) and eyes-closed (bottom rows) conditions. Purple dots represent the ASC group, and green dots represent the NT group. Solid lines indicate the average trends.

#### 3.4.1 Multiscale Sample Entropy Results

Averaging MSE across all electrodes and participants, an ANOVA confirmed increased MSE in the eyes-open condition(*F*(1, 5364) = 640.843, *p* <.001; Figure 3A). This finding aligns with prior studies (Vecchio et al., 2021), tempering our analysis pipeline. Channel-specific comparisons at individual time scales further supported these results (Figures 3B(i) and 3B(ii)).

**Figure 3.**
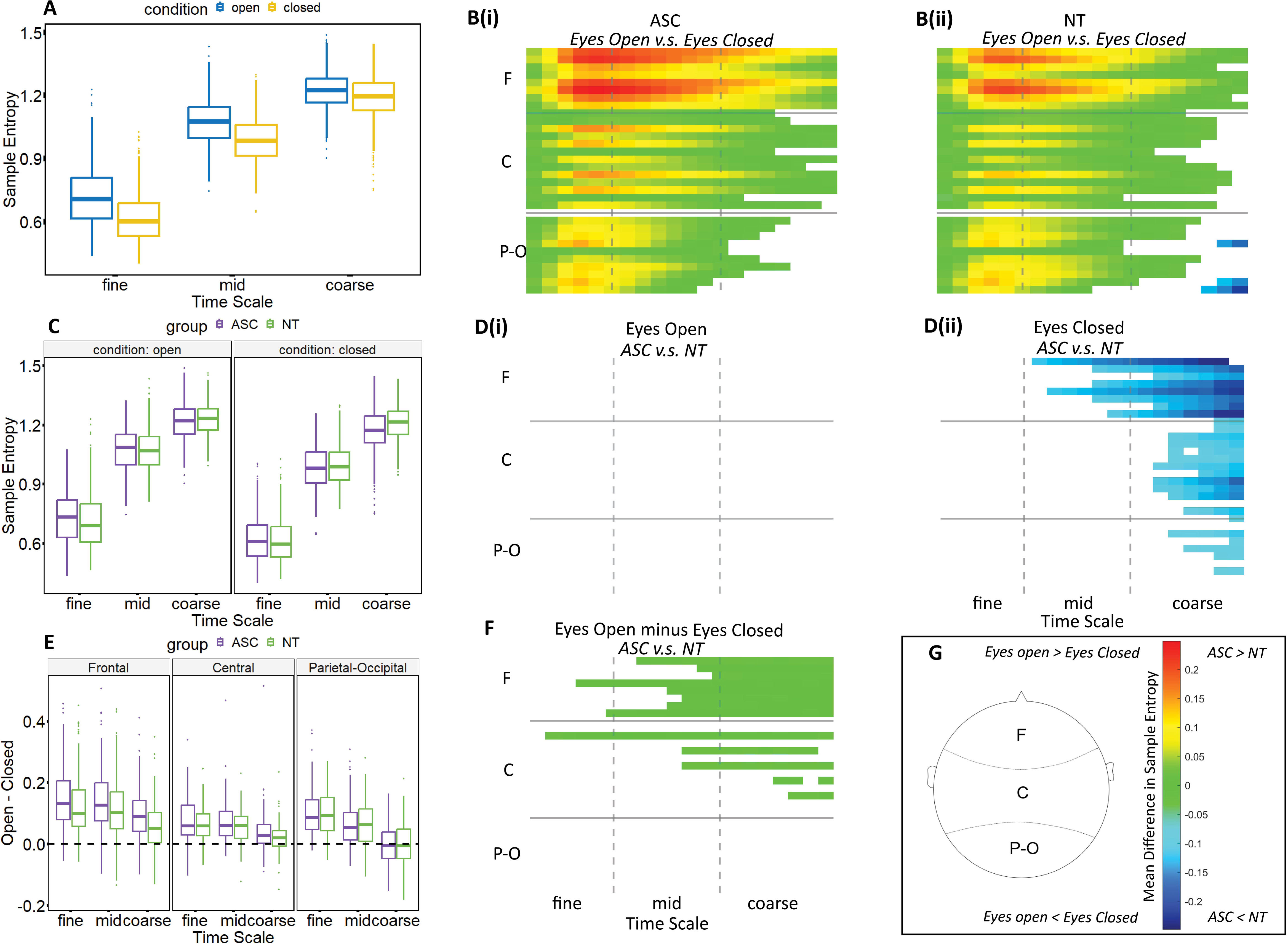
Contrasts in multiscale entropy (MSE) across diagnosis groups and eye conditions for each scalp electrode and time scale. F – frontal; C – central; PO – parieto-occipital. Plot A: Averaged MSE across subjects in eyes-open (blue) and eyes-closed (yellow) conditions for each time scale factor. Plot B(i) & B(ii): Heatmaps showing contrasts in MSE between eyes-open and eyes-closed conditions within diagnosis groups (ASC and NT). Hot colours indicate higher MSE in the specified condition, masked by significance (p < 0.05) using Threshold-Free Cluster Enhancement (TFCE) with 2000 random permutations. Plot C: Averaged MSE for ASC (purple) and NT (green) groups at eyes-open and eyes-closed conditions for each time scale factor. Plot D(i) & D(ii): Heatmaps showing contrasts in MSE between ASC and NT groups for eyes-open and eyes-closed conditions. Hot colours indicate higher MSE in the specified group, masked by significance using TFCE. Plot E: Differences in MSE between eye conditions (eyes-open minus eyes-closed) for ASC and NT groups across brain regions and time scales. Plot F: Heatmap of contrasts in MSE (eyes-open minus eyes-closed) between diagnosis groups (ASC vs NT), with significant results (TFCE, p < 0.05) shown in hot colours. Plot G: A diagram showing electrode positions with a colorbar representing the mean difference.

Fine-grained analysis using threshold-free cluster enhancement (TFCE) identified significant group differences in the eyes-closed condition (Figure 3D(ii)) and in the contrast between conditions (eyes open - eyes closed; Figures 3E, F). Heatmap figures display colour only in areas with significant group differences. These differences were concentrated in coarse time scales and frontal scalp regions. ASC participants exhibited reduced MSE compared to NT individuals during eyes-closed rest (Figure 3D(ii)) and showed larger differences between eye conditions than NT participants (Figure 3F).

#### 3.4.2 Relative Power Results

An ANOVA on data averaged across electrodes and participants revealed a significant interaction between frequency band and eye condition (*F*(5, 10728) = 542.857, p <.001; Figure 4A). Channel-specific analyses using TFCE showed decreased alpha power and increased power in other frequency bands for both groups during the eyes-open condition (Figures 4B(i) and 4B(ii)). The well documented (e.g. Berger 1929) expected increase in alpha activity during eyes-closed during rest further validated our analysis pipeline (Berger, 1929).

**Figure 4.**
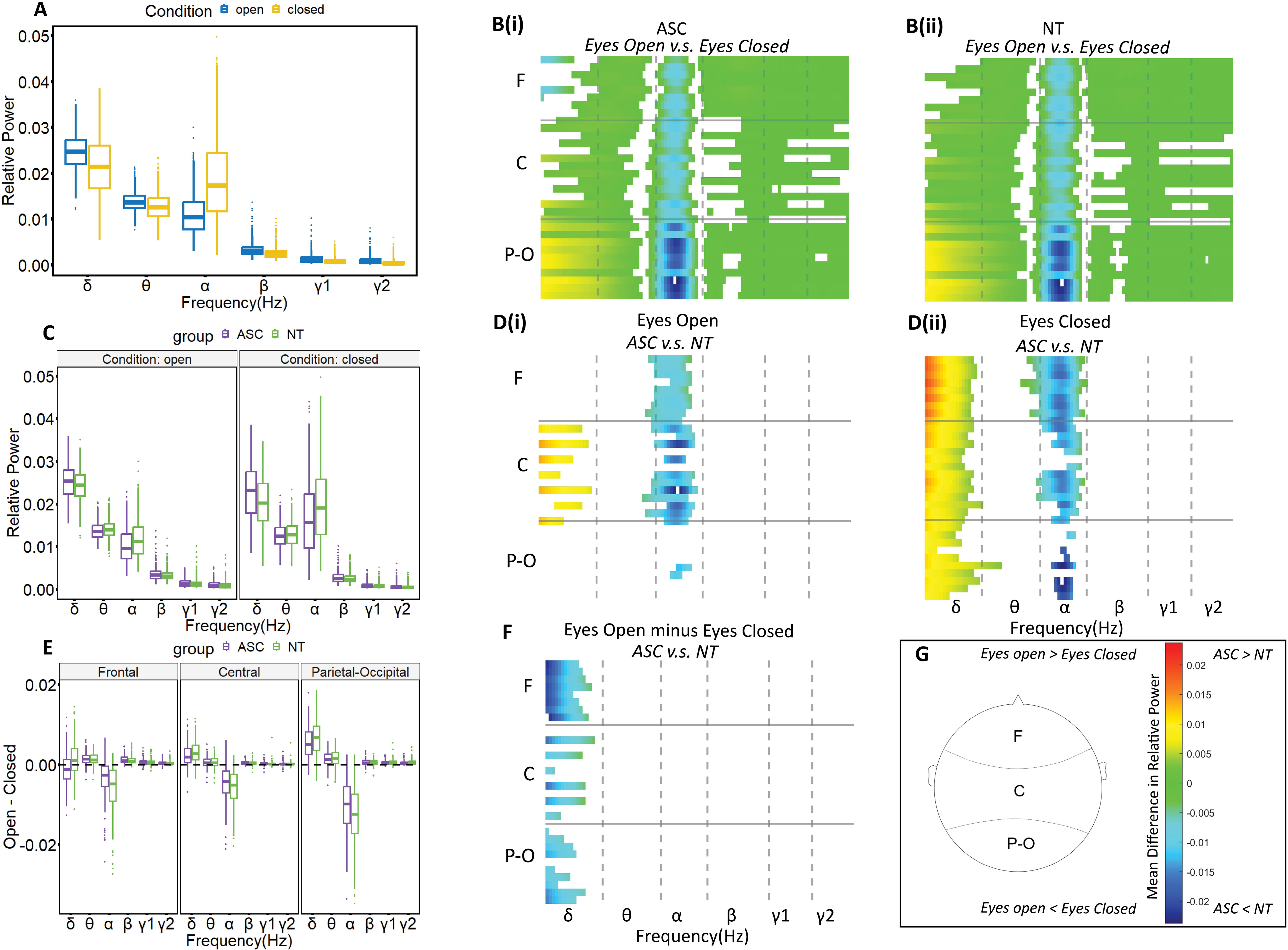
Contrasts in Relative Power across diagnosis groups and eye conditions for each scalp electrode and frequency band. F – frontal; C – central; PO – parieto-occipital. Plot A: Averaged Relative Power across subjects in eyes-open (blue) and eyes-closed (yellow) conditions for each frequency band (δ, θ, α, β, γ1, γ2). Plot B(i) & B(ii): Heatmaps showing contrasts in Relative Power between eyes-open and eyes-closed conditions within diagnosis groups (ASC and NT). Hot colours indicate higher Relative Power in the specified condition, masked by significance (p < 0.05) using Threshold-Free Cluster Enhancement (TFCE) with 2000 random permutations. Plot C: Averaged Relative Power for ASC (purple) and NT (green) groups at eyes-open and eyes-closed conditions for each frequency band. Plot D(i) & D(ii): Heatmaps showing contrasts in Relative Power between ASC and NT groups for eyes-open and eyes-closed conditions. Hot colours indicate higher Relative Power in the specified group, masked by significance using TFCE. Plot E: Differences in Relative Power between eye conditions (eyes-open minus eyes-closed) for ASC and NT groups across brain regions and frequency bands. Plot F: Heatmap of contrasts in Relative Power (eyes-open minus eyes-closed) between diagnosis groups (ASC vs NT), with significant results (TFCE, p < 0.05) shown in hot colours. Plot G: A diagram showing electrode positions with a colorbar representing the mean difference.

Comparing ASC and NT participants using TFCE, group differences were observed across all eye conditions (Figure 4D(i), 4D(ii), and 4F). Notably, ASC participants exhibited higher relative delta power and lower relative alpha power compared to the NT participants across the entire brain in the eyes-closed condition. During the eyes-open rest condition, differences were limited to frontal and central regions. In the contrast between eye conditions, group differences were only apparent in the delta frequency band, with ASC individuals exhibiting smaller changes in delta power between eye conditions than NT individuals (Figure 4F).

## 4. Discussion

In a large sample of 300 participants (126 ASC), we analyzed group differences in rs-EEG under eyes-open and eyes-closed conditions, as well as differences between these states. Our exploratory analysis of 726 rs-EEG variables revealed that meaningful group differences (η^2^*_partial_* >.035) were most prevalent for relative power and multiscale sample entropy (MSE) during the eyes closed condition. These results were confirmed with split-test validation. Fine-grained analysis using cluster-based permutation testing revealed specific frequency and topographic patterns. These analyses revealed a converging set of results showing that ASC participants in the eyes-closed condition had increased frontal coarse MSE, elevated frontal relative delta power, and reduced frontal relative alpha power.

Our findings highlight that diagnosis group differences vary by eye condition. Of the 105 significant rs-EEG variables ( η^2^*_partial_* >.035), 57 were detected in the eyes-closed condition.

Previous meta-analyses have similarly reported that differences between ASC and NT groups vary by eye-condition (Neo et al. 2023; Newson and Thiagarajan 2018). Neo et al. (2023) showed that the resting-state paradigm (eyes-open vs. eyes-closed) significantly moderates effect sizes for specific EEG frequency bands, including absolute beta power and relative delta and alpha power. Our study extends these results with a within-subjects comparison. The greater detection of group differences in the eyes-closed condition may result from its reliance on internal brain processes (Marx et al., 2003), consistent with the tendency of individuals with ASC to focus more on internal cues (Noel et al., 2018). Supporting this, recent research found that alpha microstates were reduced in stability, frequency, and engagement in ASC during eyes-closed rest, indicating disruptions in the default mode network and regions involved in self-referential thinking (Das et al. 2024). The eyes-closed condition also amplifies dominant resting-state rhythms, such as alpha oscillations, which are closely linked to cortical idling and neural inhibition (Pfurtscheller et al. 1996; Klimesch et al. 2007). These rhythms are particularly relevant to neurodevelopmental conditions like ASC (Neo et al. 2023), and their enhanced presence in the eyes-closed state may facilitate identification of group differences.

Setting aside the effects of eye conditions, differences in MSE and power changes between individuals with ASC and NT individuals have been reported. In our study, ASC participants showed lower frontal MSE at coarse scales compared to NT individuals, consistent with prior research. Hadoush et al. (2019) reported reduced MSE in the parietal and right frontal areas in children with severe ASC, and Bosl et al. (2011) observed decreased MSE in the frontal regions of infants at high risk for autism. Reduced MSE at coarse scales may indicate decreased long-range coordination (Schnitzler & Gross, 2005; von Stein & Sarnthein, 2000). Additionally, autistic participants exhibited increased delta relative power and reduced alpha relative power compared to NT participants, consistent with the U-shaped power pattern observed in ASC. This pattern is characterised by elevated low-frequency (delta) and high-frequency power, with reduced mid-frequency (alpha) power (Wang et al., 2013; Coben et al., 2008). Decreased alpha power, based on the alpha-inhibition hypothesis, may reflect lower brain inhibition (Jensen & Mazaheri, 2010), while increased delta power suggests a dominance of local brain processes over long-range connections (Nunez, 1995; Nunez & Srinivasan, 2006).

Our study did not replicate previously reported differences in connectivity, PAC, PAF, or 1/f slope observed in prior research (Duffy & Als, 2012; Liloia et al., 2024; Shou et al., 2017; Port et al., 2019; Manyukhina et al., 2022; Edgar et al., 2019; Dickinson et al., 2018). Three reasons may explain these null findings. First, as we emphasise above, brain state can influence the detection of group differences, so our consideration of eye condition may have partially controlled for state differences. Second, our study utilised a larger-than-average dataset. Larger sample sizes, as shown in our previous work and other studies, often reveal fewer rsEEG differences between ASC and NT participants as sample size rises, possibly due to heterogeneity between autistic individuals (Dede et al., 2023;Garcés et al., 2022; Kliemann et al., 2024). Third, as emphasised in our previous report (Dede et al. 2023), heterogeneity can lead to the detection of effects that do not survive validation methods. For example, Figure 5A displays a histogram of raw alpha power in both groups over left-frontal scalp locations during the eyes closed condition. Visual inspection reveals that the ranges of the two groups are completely overlapping.While this variable had a high effect size (0.047), its reliability was low (0.313). By contrast, Figure 5B displays group histograms for relative delta power over the same region at the eyes closed condition. Visual inspection reveals that the ranges of the two groups do not completely overlap. This variable yielded both a high effect size (0.068) and high reliability (0.813). This comparison highlights that relying solely on effect size or p-value without doing validation analysis can be misleading.

**Figure 5.**
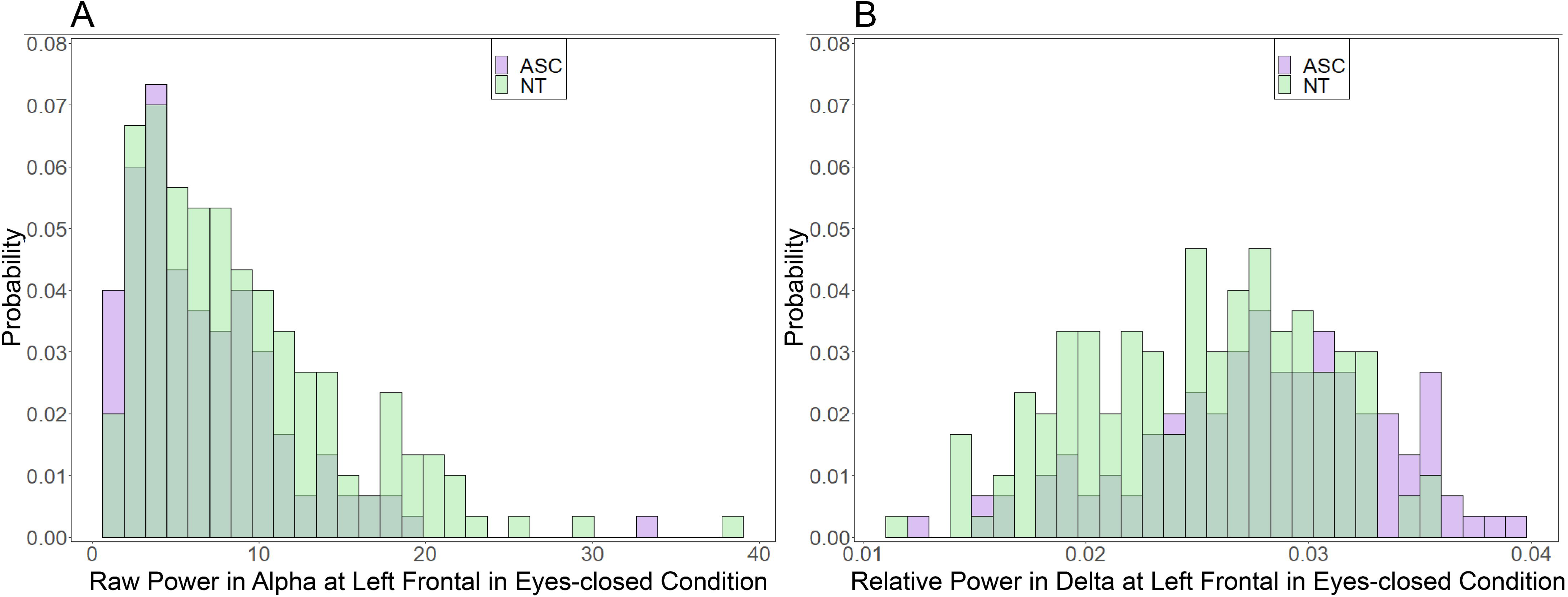
Group-level histograms demonstrate the importance of reliability in interpreting EEG group differences. A: Histogram of raw alpha power. Visual inspection reveals significant overlap in the distributions of ASC and NT groups, indicating low reliability of the observed group difference despite a high effect size. Effect size: 0.047. Reliability index: 0.313. Descriptive statistics (group: mean (SD)): ASC: 6.79 (4.83); NT: 9.00 (6.15).B: Histogram of relative delta power. The distributions for ASC and NT groups show minimal overlap, resulting in both a high effect size and high reliability for this variable. Effect size: 0.068. Reliability index: 0.813. Descriptive statistics (group: mean (SD)): ASC: 0.0280 (0.00545); NT: 0.0250 (0.00527).

Cross-validation is crucial for enhancing statistical rigour and reproducibility, particularly in studies where findings need to be validated across independent datasets. For example, a previous report using the same data analysed here reported that alpha power was lower in ASC participants during eyes open rest (Neuhaus et al., 2021); our analysis also arrived at this finding, however this result did not survive cross validation. More generally, even when effects seem large and machine learning approaches are capable of categorising between diagnostic groups with high accuracy, chance performance can be revealed when categorisation is attempted on independent datasets, emphasising the importance of cross-validation methods (Chekroud et al., 2024).

Lastly, it is important to note that although our study revealed some statistically robust effects at the group level, the size of these effects was not large. As can be seen in Figure 5, there was substantial overlap between ASC and NT individuals. This fits with the view that ASC is a heterogeneous condition which may not be fully delineated by any single physiological difference between ASC and NT individuals (Dede et al., 2023). Future research should combine control of brain state, as we have done here, with more fine-grained consideration of particular features of the autism phenotype in line with the RDOC approach (Volkmar 2013), or within homogeneous subgroups of autistic individuals (Molloy & Gallagher, 2022). In a similar conceptual vein, it may also be fruitful to consider how changes in brain state vary between groups defined by differences in genotype. For example, grouping participants by genotype can predict fMRI connectivity better than mental health diagnostic labels (Moreau et al., 2023).

## Limitations

Our analysis had several limitations. First, because our analysis was exploratory, future research should aim to replicate our findings. Second, we focused on univariate rs-EEG data due to its accessibility and clinical relevance. We cannot rule out the possibility that clearer EEG differences between autistic and neurotypical individuals may manifest during task engagement or in multivariate analyses. Third, given that ASC is a neurodevelopmental condition and age is a strong predictor of EEG metrics, as indicated in this study, our previous study (Dede et al., 2023) and many prior reports (Lawrence et al. 2019; van Noordt and Willoughby 2021), conducting longitudinal research becomes imperative. Finally, we cannot rule out the possibility that there are further variables that could be extracted from the rs-EEG signal that may more successfully differentiate the autistic from the neurotypical sample. However, our analysis approach was comprehensive and included indices that represent fundamental features of neural dynamics, therefore it would be somewhat surprising for a novel measure to demonstrate meaningful group differences.

## Conclusions

In sum, there are three key takeaways from this work. First, eye condition impacts the detection of group differences between ASC and NT individuals. We found more prominent differences in the eyes-closed condition than the eyes-open condition. Second, we found that robust differences between ASC and NT individuals were most prevalent in measures of relative spectral power and coarse MSE over frontal scalp locations. Third, even with the added control over brain state provided by consideration of eye condition, group differences were few and modest, which may be caused by the heterogeneity between individuals.

These findings suggest a ‘both-and’ approach for future research. On one hand, researchers may benefit from more refined participant selection, such as subtyping individuals based on specific genetic or phenotypic variants of autism, as recommended in our previous study (Dede et al., 2023). On the other hand, careful control of state variables, as demonstrated through analysis of eye conditions in this study, should also be considered. This dual approach could provide a more comprehensive understanding of the interaction between state and trait influences on neural dynamics and aid in the detection of reliable group differences.

## Abbreviations

ASC: Autism Spectrum Condition
NT: neurotypical
rs-EEG: resting-state electroencephalography
MSE: multiscale entropy

## Acknowledgements

We sincerely thank the Autism Center for Excellence (ACE) team for granting access to their dataset and the National Database for Autism Research (NDAR) for hosting the data. We also acknowledge the University of Sheffield for providing high-performance computing resources essential for data analysis.

